# Coiled-coil domain kinking controls laminin-332 cleavage by elastase

**DOI:** 10.64898/2026.02.08.704728

**Authors:** Lucky Akter, Romain Amyot, Robert Großmann, Carsten Beta, Holger Flechsig, Clemens M. Franz

## Abstract

Laminins are trimeric glycoproteins and essential components of basement membranes, where they provide structural support and mediate cell-matrix adhesion. Their α-, β-, and γ-chains combine into a flexible trimeric coiled-coil domain forming the laminin long arm, which links the N-terminal short arms and the C-terminal globular domains. However, ultrastructural insight into the laminin coiled-coil domain remains limited, and the function of coiled-coil flexibility remains unclear. Using high-speed atomic force microscopy (HS-AFM) imaging, we previously investigated the ultrastructure of laminin-332 and observed dynamic kinking around a putative molecular hinge in the center of the coiled-coil domain. In this study, we integrate HS-AFM time-lapse imaging with AlphaFold structure prediction and normal-mode flexible fitting (NMFF) to generate dynamic models of the laminin-332 hinge during coiled-coil domain kinking, providing atomic-scale insight into the underlying molecular rearrangements. Furthermore, we use HS-AFM to visualize, in real time, the digestion of individual laminin-332 molecules by pancreatic elastase and show that coiled-coil kinking directs elastase cleavage to the hinge site, thereby ensuring reliable generation of the elastase 8 (E8) fragment containing the integrin binding site. In contrast, laminin molecules with extended coiled-coil conformations are cleaved at arbitrary sites, resulting in complete coiled-coil removal and degradation. Our results thus identify a novel conformational control mechanism directing proteolytic processing of laminin-332 through defined coiled-coil kinking. Besides expanding the understanding of dynamic laminin coiled-coil function, the gained ultrastructural insight into trimeric coiled-coil bending may aid the rational design of flexible coiled-coil modules for future protein engineering applications.

## Introduction

Laminins are large glycoproteins with important roles in organizing basement membranes, thin extracellular matrix sheets that provide support for epithelial and endothelial cells and separate them from the underlying connective tissue^1, 2^. Within basement membranes, laminins form supramolecular networks through self-aggregation^3^ or via interactions with other basement membrane constituents, like collagen IV and nidogen, and proteoglycans such as perlecan and agrin^4^. In addition to their scaffolding function, laminins contain binding sites for integrins and other cell surface receptors and modulate cell signaling pathways during embryogenesis, differentiation, neurite outgrowth, and angiogenesis^5^, as well as cancer metastasis^6, 7^. Laminin mutations that compromise basement membrane integrity are also associated with a range of inherited disorders, including congenital muscular dystrophy, cardiomyopathies, and epidermolysis bullosa^8^, underlining the importance of laminins for basement membrane and overall tissue function.

Laminins are heterotrimers composed of one α-, one β-, and one γ-chain. In mammals, there are 16 isoforms with tissue-specific expression profiles and functions that arise from different combinations of five α-, three β-, and three γ-chains^9^. The N-termini of the three laminin chains form the laminin short arms and contain various protein interaction sites, including LN domains for facilitating laminin self-aggregation^10, 11^, and nidogen^12^, which in turn facilitates binding to collagen IV and the heparan sulfate proteoglycan perlecan^13^. The C-terminal sections of the three laminin chains combine into the laminin long arm by forming an extended triple coiled-coil domain spanning ∼600 amino acid residues. Coiled-coil domains are common protein-protein interaction domains in which two or more α-helices are held together by hydrophobic interactions and coiled into a superhelix, which requires a repeating seven amino acid (heptad) pattern with hydrophobic residues in the first and fourth position of the heptad^14^. Tight packaging of the hydrophobic amino acids into the core provides thermodynamic driving force for coiled-coil self-assembly. The laminin coiled-coil domain is further stabilized by disulfide bridges, while ionic interactions within the coiled-coil domain govern the specificity of laminin chain assembly^15, 16^. Laminin α-chains extend beyond the C-terminus of the coiled-coil domain and terminate in a cluster of five LG domains (LG1-5), which harbor binding site for integrins and other glycoproteins and proteoglycans, such as α-dystroglycan^17^ and syndecan-1^18^. In contrast to the N- and C-terminal domains, only few protein interactions have been described for the laminin coiled-coil^19^, which may primarily serve as a molecular spacer between the terminal protein binding sites^20^.

Combining a specific subset of α-, β-, and γ-chains generates laminin isoforms tailored to different tissue requirements. For instance, the laminin-332 isoform (α3β3γ2) is expressed in skin where it is essential for anchoring keratinocytes to the underlying dermis at the dermal–epidermal junction, a specialized basement membrane structure ensuring skin integrity and resistance against external mechanical forces^21^. Loss-of-function mutations of any of the three laminin-332 chains (α3, β3, or γ2) drastically decrease dermal-epidermal adhesion and give rise to junctional epidermolysis bullosa, a lethal condition characterized by extreme skin fragility and blistering^22^. The tissue-specific functions of laminin-332 are further modulated by extensive proteolytic processing^23^, such as trimming of the N-termini of the α3- and γ2-chains^24, 25^ and cleavage of the linker between the C-terminal LG3 and LG4 domains^26^, leading to the release of the LG4-5 module. Laminin-332 is furthermore cleaved by matrix metalloproteinases (MMPs) and neutrophil elastase, releasing promigratory fragments^27, 28^. Moreover, the laminin α3 chain can be expressed as a shorter splice variant A (α3A) or the long B variant (α3B)^29^. Consequently, the specific functions of the laminin-332 isoform arise from the combined effect of laminin chain selection, transcriptional processing, and posttranslational proteolytic processing, and understanding the precise influence of these modifications on laminin-332 function remains challenging.

Given their essential roles in tissue homeostasis and disease, elucidating the structure–function relationships of laminins has been a significant focus of biomedical research^30, 31^. X-ray crystallography and cryo-electron microscopy (EM) studies have generated a large collection of N-terminal and C-terminal laminin substructures, including the ternary LN complex underlying laminin network assembly^11^ and the C-terminal LG domains mediating integrin binding^32, 33^. In contrast, comparable atomic-scale insight into coiled-coil domain structure is still missing, because its flexible nature impedes crystallization or single particle averaging for electron microscopy. The understanding of laminin long arm structure is therefore still largely based on early EM^34, 35^ and AFM^36^ images and on computational approaches^37^. The pioneering EM images demonstrated the characteristic cross-shaped structure of laminins and revealed diverse long arm shapes, suggesting a high degree of flexibility of the central coiled-coil domain^38^. However, these images represent only selected static snapshots rather than a full set of conformational states of the coiled-coil domain. Understanding the full range of coiled-coil domain conformations and their potential importance for laminin function therefore requires imaging methods capable of monitoring dynamic conformations with high temporal and spatial resolution.

High-speed atomic force microscopy (HS-AFM) operates under physiological conditions and can visualize molecular rearrangement processes with subsecond and nanometer resolution^39^. We previously employed HS-AFM to investigate laminin ultrastructure and observed dynamic kinking of the laminin-332 coiled-coil domain and identified a putative molecular hinge located in the center of the coiled-coil domain^40^, providing new isoform-specific insight into laminin coiled-coil domain behavior. However, HS-AFM images of proteins have a spatial resolution typically limited to ∼1-2 nm and therefore they cannot reveal structural rearrangements with atomic resolution. To overcome these previous limitations, here we have combined HS-AFM imaging, AlphaFold-based structure prediction^41^, and a normal mode flexible fitting (NMFF) algorithm^42, 43^ to generate atomistic models describing the molecular changes occurring during reversible laminin-332 coiled-coil domain kinking. Furthermore, using HS-AFM we visualize cleavage of individual laminin-332 molecules by elastase and show that coiled-coil domain kinking reliably directs elastase cutting to the hinge region, ensuring the production of the biologically active elastase 8 (E8) fragment. Our results thus provide a first ultrastructural view into a dynamic coiled-coil hinge and demonstrate a novel function for laminin-332 coiled-coil domain kinking in regulating its susceptibility to protease digestion.

## Materials and Methods

### Sample preparation and HS-AFM observation

Human recombinant laminin-3B32 (LN332-0202) was purchased from Biolamina (biolamina.com). A stock solution (100 µg/ml) was aliquoted and stored at-80°C. Recombinant human laminin-332 E8 fragment (“iMatrix-332) was purchased from Matrixome (matrixome.co.jp), aliquoted (0.5 mg/ml) and stored at-80°C. Lyophilized Porcine pancreatic elastase (CAS 39445-21-1) was purchased from Merk (#324682) and stored at-20°C or reconstituted as a working solution of 6.4 mg/ml in 150 mM NaCl, 50 mM MES, pH6 and stored at 4°C for use within 2 weeks. Just before HS-AFM imaging, the full length laminin-332 or laminin-332 E8 fragment stock solution was diluted to 1 µg/ml in laminin imaging buffer (150 mM NaCl, 50 mM Tris, pH 7.5). A muscovite mica disc (0.1 mm thickness, 2 mm diameter) attached to a sample glass stage (2 mm diameter, 2 mm height) with cyanoacrylate glue was then cleaved and immediately incubated with 0.01% (3-aminopropyl) triethoxysilane (APTES) in ddH_2_0 and incubated for 3 minutes, followed by 5 washes each with ddH_2_0 and laminin imaging buffer. Afterwards, 2 µl laminin solution was applied to the APTES-coated mica disc and incubated for 10 min. After five washes with imaging buffer, the sample stage was inserted into the chamber of a laboratory-built HS-AFM with a narrow-range scanner^44^. Images were acquired using BL-AC10DS-A2 cantilevers (Olympus) with a spring constant of 0.1 N/m and a resonance frequency of ∼0.4 MHz in water. The cantilevers carried an amorphous carbon tip fabricated on the original tip by electron beam deposition (EBD) using a Verios 460 scanning electron microscope (FEI). The length of the additional AFM tip was ∼300-500 nm, and the tip apex radii typically ranged between ∼1 and 4 nm. Tips were further sharpened by plasma etching. A laser beam (670 nm) was focused on the cantilever tip through a 20X objective lens to detect the cantilever deflection. The free oscillation amplitude of the cantilever was set to ∼2 nm and the set-point amplitude for the feedback control to 80-90% of the free amplitude. Feedback parameters were optimized during scanning to minimize tip-sample interaction forces. All HS-AFM experiments were performed at room temperature. To study elastase digestion of laminin-332, 10 µl of elastase working solution (6.4 mg/ml) was added during HS-AFM imaging of laminin-332 molecules into the AFM sample chamber containing 90 µl of laminin imaging buffer.

### HS-AFM image processing and data analysis

HS-AFM raw images were processed in Gwyddion 2.66^45^ by applying mean plane subtraction, row alignment, and correcting horizontal scan lines if needed. HS-AFM movies were flattened by applying a plane or second-order polynomial surface fitting using in-house software routines developed in MATLAB (Mathworks, R2021)^46^, or the NanoLocz software^47^. Image frame drift was corrected using 2D alignment image registration features of ImageJ (www.imagej.nih.gov/ij/). Laminin coiled-coil length and kink angles were measured using ImageJ and plotted in Origin Pro 2021 (www.originlab.com).

### Determining coiled-coil kink angle distribution, kink angle potential, spring constants

A list of 209 experimentally measured coiled-coil domain kink angles {𝜑_1_, 𝜑_2_,…} revealed a distinctly bimodal distribution 𝑃_𝑠_(𝜑), reminiscent of the superposition of two Gaussians corresponding to characteristic opening angles in a kinked and an extended configuration. To obtain an analytical expression for the probability distribution, we hence inferred a two-component Gaussian mixture model

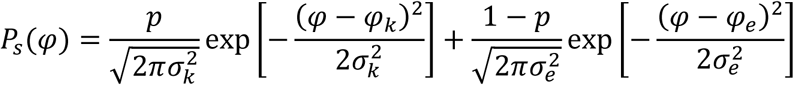

from the data employing an expectation-maximization algorithm^48^ as implemented in scikit-learn. The following mean values and standard deviations were found: 𝜑_𝑘_ = 77^∘^, 𝜎_𝑘_ = 19^∘^, 𝜑_𝑒_ = 159^∘^, 𝜎_𝑒_ = 13^∘^. The relative weight factor was estimated to be 𝑝𝑝 = 0.69, reflecting a higher probability to find small angles. In thermal equilibrium, the stationary distribution of kink angles is expected to follow a Boltzmann distribution

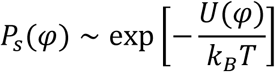

with the absolute temperature 𝑇. Using the Gaussian mixture model, the kink angle potential 𝑈(𝜑) was inferred up to an arbitrary constant 𝑈_0_ via:

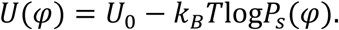

The potential energy revealed two minima corresponding to the two metastable states, with transition barrier heights of 𝑈_𝑘_ ≈ 2.73 𝑘_𝐵_𝑇 and 𝑈_𝑒_ ≈ 2.32 𝑘_𝐵_𝑇. Close to its minima, the kink angle potential is well approximated by a harmonic potential

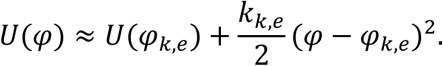

The spring constant of the potential in the two metastable states were estimated using the Boltzmann distribution and the Gaussian mixture model,

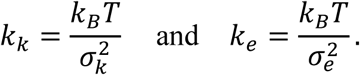

The ratio of the spring constants is determined by the widths of the respective peaks:

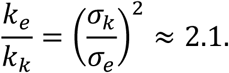

Thus, the kink angle potential is stiffer in the extended configuration, but also shallower, than in the kinked configuration.

### Life-time estimations of the extended and kinked conformations

The time series of angle measurements were typically not long enough to estimate the transition rates (or, equivalently, the mean live time of the two characteristic states) directly. Instead, we relied on inferring a stochastic model from the data that allows for longer simulations, which in turn was used to estimate the respective characteristic transition times. The dynamics of the kink angle 𝜑𝜑(𝑡𝑡) is assumed to follow overdamped Langevin dynamics

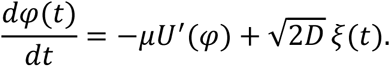

The dynamics is composed of two parts: a deterministic force 𝑓(𝜑) = −𝑈^′^(𝜑) = −𝑑𝑈(𝜑)/𝑑𝜑 which derives from a potential 𝑈(𝜑) (see *kink angle potential* above), and a fluctuating force 𝜉(𝑡) that is described by Gaussian white noise with zero mean, ⟨𝜉(𝑡)⟩ = 0, and temporal 𝛿𝛿-correlations: ⟨𝜉(𝑡)𝜉(𝑡^′^)⟩ = 𝛿(𝑡 − 𝑡^′^). In thermodynamic equilibrium, the fluctuation-dissipation relation 𝐷𝐷 = 𝜇𝜇𝑘𝑘_𝐵𝐵_𝑇𝑇 holds, ensuring the stationary kink angle distribution to be of Boltzmann-type:

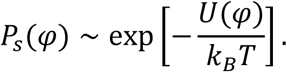

The relative potential 𝑈(𝜑)/(𝑘_𝐵_𝑇) can be obtained by Boltzmann inversion, 𝑈(𝜑)/(𝑘_𝐵_𝑇) = const. − log𝑃_𝑠_(𝜑), in which the stationary distribution 𝑃_𝑠_(𝜑) is estimated from the time series using a two-component Gaussian mixture model as described above. The Langevin dynamics can thus be rewritten as follows:

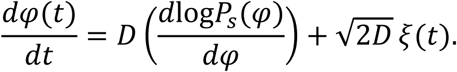

The parameter 𝐷𝐷, setting the timescale of the dynamics, can be estimated from the times series by likelihood maximization (see Supplementary Information) and estimated as 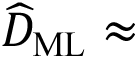 0.12 rad^2^/s. Simulating the Langevin dynamics numerically, we estimate the lifetimes of the two characteristic configurations as follows: the dynamics is initially in one of the two states by drawing an initial angle at random from the respective Gaussian distribution, and the time until the angle 𝜑 passes through the energy maximum (𝜑_𝑚_ = 126^∘^, cf. Fig. 2F) is measured; this process is repeated 10^4^times. The lifetimes are exponentially distributed, with the following mean waiting times for the respective transitions: 𝜏_𝑘→𝑒_ ≈ 12.7 s, 𝜏_𝑒→𝑘_ ≈ 2.6 s.

**Figure 1.**
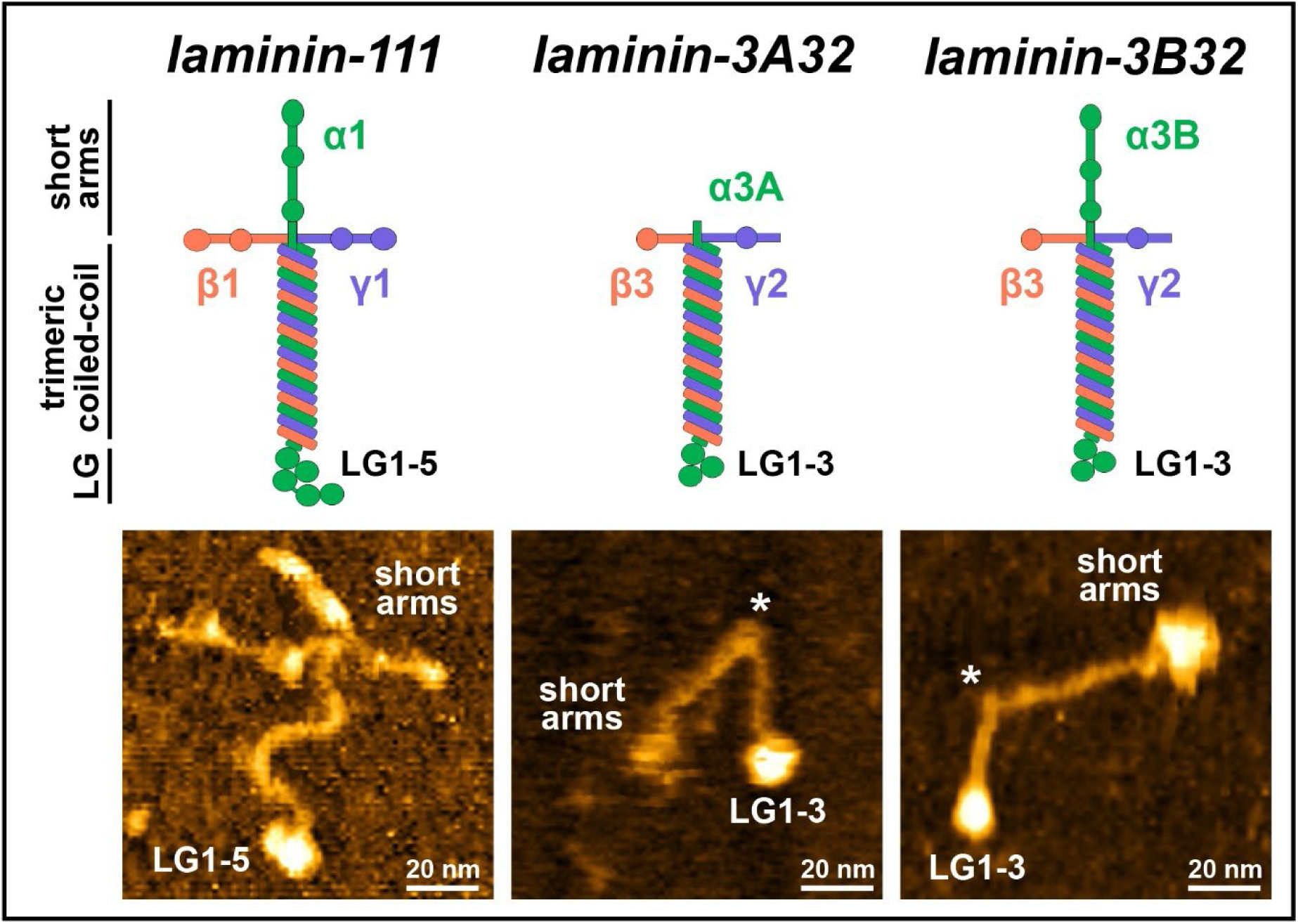
Laminin-332 isoforms display a kinked trimeric coiled-coil domain. Schematic depiction of laminin-111, laminin-3A32 and laminin-3B32 (upper row) and corresponding AFM images (lower row). Laminin molecules were immobilized on APTES-functionalized mica surfaces. Asterisks denote the position of a molecular hinge within the trimeric coiled-coil domain. The full range of the height scale is 8 nm.

**Figure 2.**
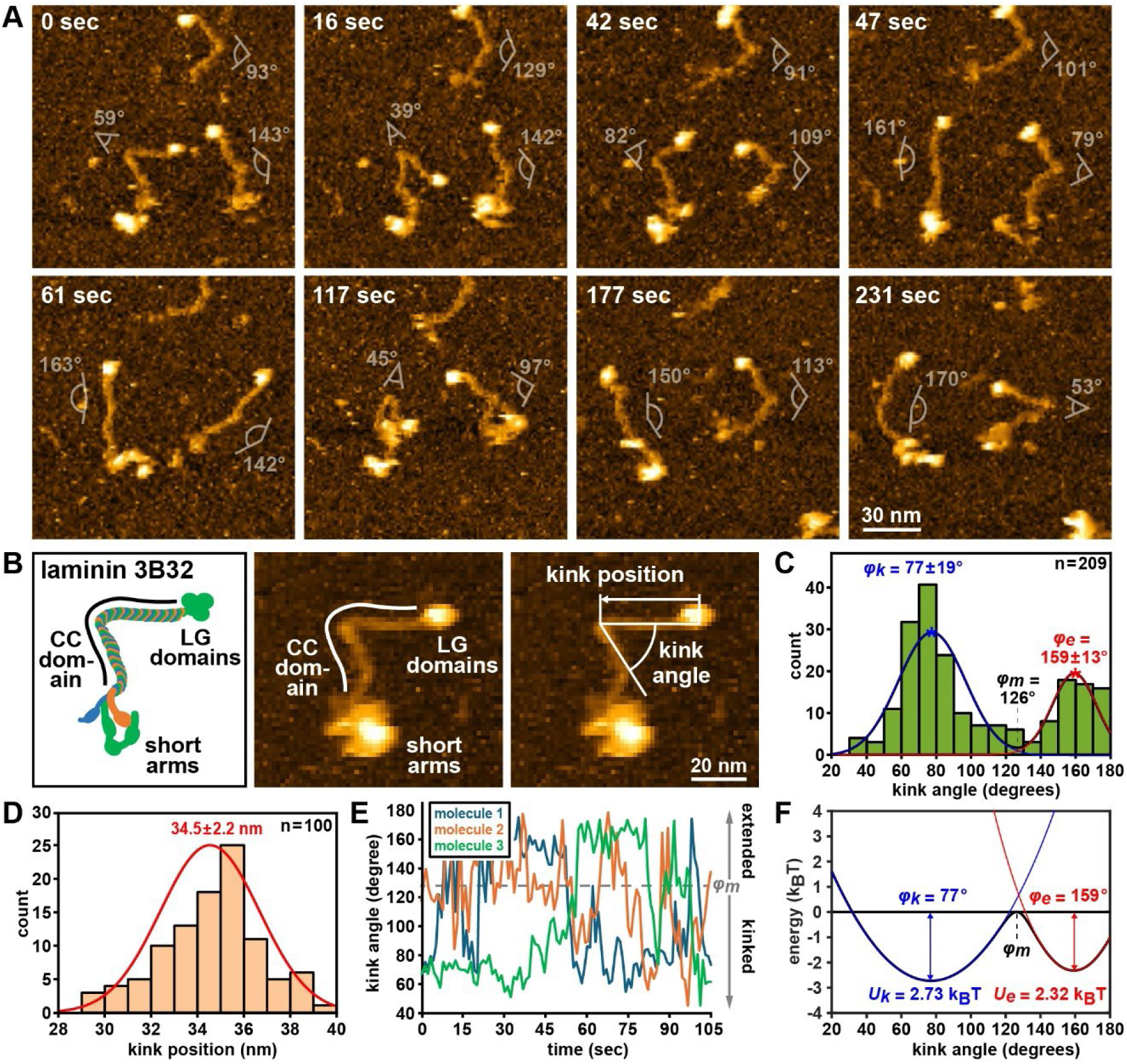
Dynamic coiled-coil kinking in laminin-332. **(A)** Stills from a HS-AFM timelapse series of laminin-332 molecules immobilized on APTES-coated mica. Coiled-coil (CC) domain kink angles are indicated next to each molecule. **(B)** Schematic depiction of laminin-3B32 (left panel) and illustration of kink angle and position analysis. **(C)** Kink angle distribution obtained from 209 measurements of 10 different molecules. *φ_k_* denotes the angle of the kinked and *φ_e_* of the extended conformation (mean ± SD), *φ_m_* is the angle at the local minimum. **(D)** Distribution of the kink position (mean ± SD) measured from the C-terminal end of the coiled-coil. **(E)** Kink angle time traces of three molecules imaged over 105 sec by HS-AFM, *φ_m_* denotes the transition angle between kinked and extended conformations. **(F)** Energy profile obtained through Boltzmann inversion of the bimodal kink angle distribution in **(C)**, *U_k_* denotes the energy barrier height for transition from the kinked into the extended conformation, *U_e_* the barrier height for transition from the extended into the kinked conformation.

### Structure prediction of the laminin coiled-coiled/elastase complex

Structure predictions of the central coiled-coil domain stretch containing the hinge region of Human laminin-332 were performed using the AlphaFold server powered by AlphaFold3^41^ and the following amino acid sequences: LAMA3_HUMAN (Q16787) Lys^2050^ – Asp^2300^, LAMB3_HUMAN (Q13751) Gly^827^ – Gly^1077^, and LAMC2_HUMAN (Q13753) Arg^843^ – Val^1093^. Models of the laminin/elastase complex additionally included the following sequence of porcine pancreatic elastase CELA1_PIG (P00772) Val^27^ – Asn^266^, missing the N-terminal propeptide sequence. The AlphaFold prediction of porcine pancreatic elastase was also compared to the human equivalent CELA1_HUMAN (Q9UNI1) Val19 – Asn^258^, again excluding the propeptide. The elastase/laminin complex shown in Fig. 6D was optimized by molecular docking using HDock^49^. The AlphaFold3-generated molecular structures (rank 1 models) were visualized in space-fill or ribbon plots in ChimeraX^50^. The Alphafold3-generated model and an X-ray crystal structure of porcine pancreatic elastase (3est) were overlaid in ChimeraX, and structure similarity was determined in terms of the root mean square deviation (RMSD) of alpha-carbon atom positions for the common set of amino acid residues.

### AFM image simulation and normal-mode flexible fitting AFM

Pseudo AFM images of the laminin coiled-coil/elastase complex were generated using the AlphaFold3-generated model as input and the BioAFMviewer software^51^, which simulates AFM scanning based on the non-elastic collisions of a rigid cone-shaped tip model with the rigid van-der Waals atomistic structure. A scan step of 1 nm was used, and the tip shape parameters were 0.8 nm for the tip probe sphere radius and 5° for the cone half angle. For molecular fitting of the AlphaFold3-generated model into experimental HS-AFM topographies, we employed the normal mode flexible fitting AFM (NMFF-AFM) method^43^ implemented in the BioAFMviewer software interface^51^. The kinked structure was chosen as the initial conformation. The region for fitting was manually selected on the raw HS-AFM topographic images processed by applying automatic tilt correction of the AFM stage and a Gaussian filter with 1 nm standard deviation. The tip shape parameters were R = 3 nm for the probe radius and ϑ = 10° for the cone half angle. Flexible fitting was performed with the following set of parameters: elastic network cutoff distance 8 Å, number of normal modes 15, RMSD of structural deformation 0.5 Å, image correlation coefficient for image similarity scoring.

### Protein sequence alignment

To identify putative elastase cutting sites in laminin-332, the α-, β-, and γ-chain coiled-coil domain sequences of Mouse laminin-111 and Human laminin-332 were aligned using Protein BLAST (https://www.ncbi.nlm.nih.gov/), and the LAMA3_HUMAN (Q16787), LAMB3_HUMAN (Q13751), LAMC2_HUMAN (Q13753) and LAMA1_MOUSE (P19137), LAMB1_MOUSE (P02469), LAMC1_MOUSE (P02468) amino acid sequences.

Human laminin-332 amino acid residues corresponding to the previously established pancreatic elastase cutting sites (P1 positions) in Mouse laminin-111^52^ were then identified in the aligned sequence.

## Results

### The laminin-332 coiled-coil domain cycles spontaneously between extended and kinked conformations

We previously imaged different laminin isoforms by HS-AFM and observed dynamic cycling of the laminin-3A32 isoform between straight and bent conformations around a central molecular hinge within the laminin coiled-coil domain, while laminin-111 and other isoforms display an S-shaped, stable morphology^40^. Here we characterize dynamic coiled-coil kinking of laminin-332 in detail and explore potential biological implications of coiled-coil kinking. In these experiments we used the laminin-3B32 isoform (from here on referred to as laminin-332 for simplicity), which features truncated β- and γ-chain short arms^23^. The trimmed β- and γ-chain short arms typically bundled together with the α-chain short arm into a compact, globular structure at the laminin N-terminus, which could be easily distinguished from the smaller LG1-3 domain cluster at the laminin C-terminus (Fig.1 and Suppl. Fig. S1). This extended dumbbell shape characterized by two globular structures connected via a straight or kinked coiled-coil domain closely resembled previous rotary shadow images of laminin-332^35^. We first imaged dynamic cycling between kinked and extended conformations of laminin-332 by time-lapse HS-AFM and determined kink positions and angles from the obtained image series (Fig. 2A and B, and Movie 1). These experiments revealed a bimodal kink angle distribution ranging from 30**°** to 180**°** with peaks at 77**°** (kinked conformation) and 159**°** (extended conformation) and a kink position 34.5 ± 2.2 nm upstream of the C-terminal coiled-coil end (Fig. 2C and D). Thus, laminin-332 perpetually cycles between a favored kinked conformation (69% share) and a rarer extended conformation (31% share), with a transition angle of ∼126° between the two metastable states. Trough Boltzmann inversion of the bimodal kink angle distribution, we constructed an energy profile demonstrating barriers heights of 2.73 k_B_T (transition kinked to extended) and 2.32 k_B_T (extended to kinked, Fig. 2F). Energy barrier heights on the order of the thermal energy k_B_T indicate that reversible transitions between kinked and extended coiled-coil conformations should be easily facilitated by thermal fluctuations. In agreement, kink angle plots over time compiled from HS-AFM time-lapse movies demonstrated rapid cycling between the different conformational states (Fig. 2E), with mean dwell times of 12.7 sec in the kinked conformation (kink angle 𝜑 < 126°) and 2.6 sec in the extended conformation (kink angle 𝜑 > 126°; see Materials and Methods). We also compared the spring constant *k* of the kink angle potential (𝜑) ≈ 0.5𝑘𝜑^2^ + const. of the two metastable states and obtained a ratio for *k*_kinked_/*k*_extended_ of ∼0.48:1, indicating roughly two-fold softening of the potential in the kinked conformation.

### Revealing dynamic hinge architecture by flexible fitting of AlphaFold-generated coiled-coil models into resolution-limited AFM topographies

Rapid cycling between kinked and extended conformations around a defined flexure point indicated the presence of a molecular hinge facilitating reversible coiled-coil bending within laminin-332, as previously suggested^40^. To further characterize the dynamic molecular hinge architecture, we applied a recently developed normal-mode flexible fitting (NMFF) approach for fitting molecular structures as elastic networks into resolution-limited AFM images^42^. This fitting approach overcomes the inherent spatial resolution limit of AFM (∼1-2 nm in *x*,*y*) but requires atomic models of the imaged molecules as structural templates. We previously showed that AlphaFold can reliably predict short segments of trimeric laminin coiled-coil domains^40^. Here, we modelled a 250 amino acid segment from the central region of laminin-332 containing the putative hinge region in AlphaFold (Fig. 3A). The predicted structure displayed the expected rod-like structure, proper trimeric coiled-coil packing and a central bend with an angle of ∼140**°**. Prediction confidence was high (pLDDT score >70%) to very high (pLDDT score >90%) throughout the structure, except at the coiled-coil termini containing the loose laminin chain ends (Fig. 3B). Interestingly, the bend region containing the putative hinge also showed reduced prediction confidence, suggesting greater flexibility in this area. The predicted coiled-coil segment was then fitted by NMFF into topographic AFM images depicting the laminin-332 coiled-coil in different bending conformations (Fig. 3C and D and Movie 2). Before starting the iterative fitting protocol, the AlphaFold structure and the topographic AFM images were manually overlayed to ensure proper N-/C-terminal orientation, while the central bend served as a positional marker to align the model with the hinge region in the AFM image. Flexible fitting readily transformed the slightly bent (∼140°)

**Figure 3.**
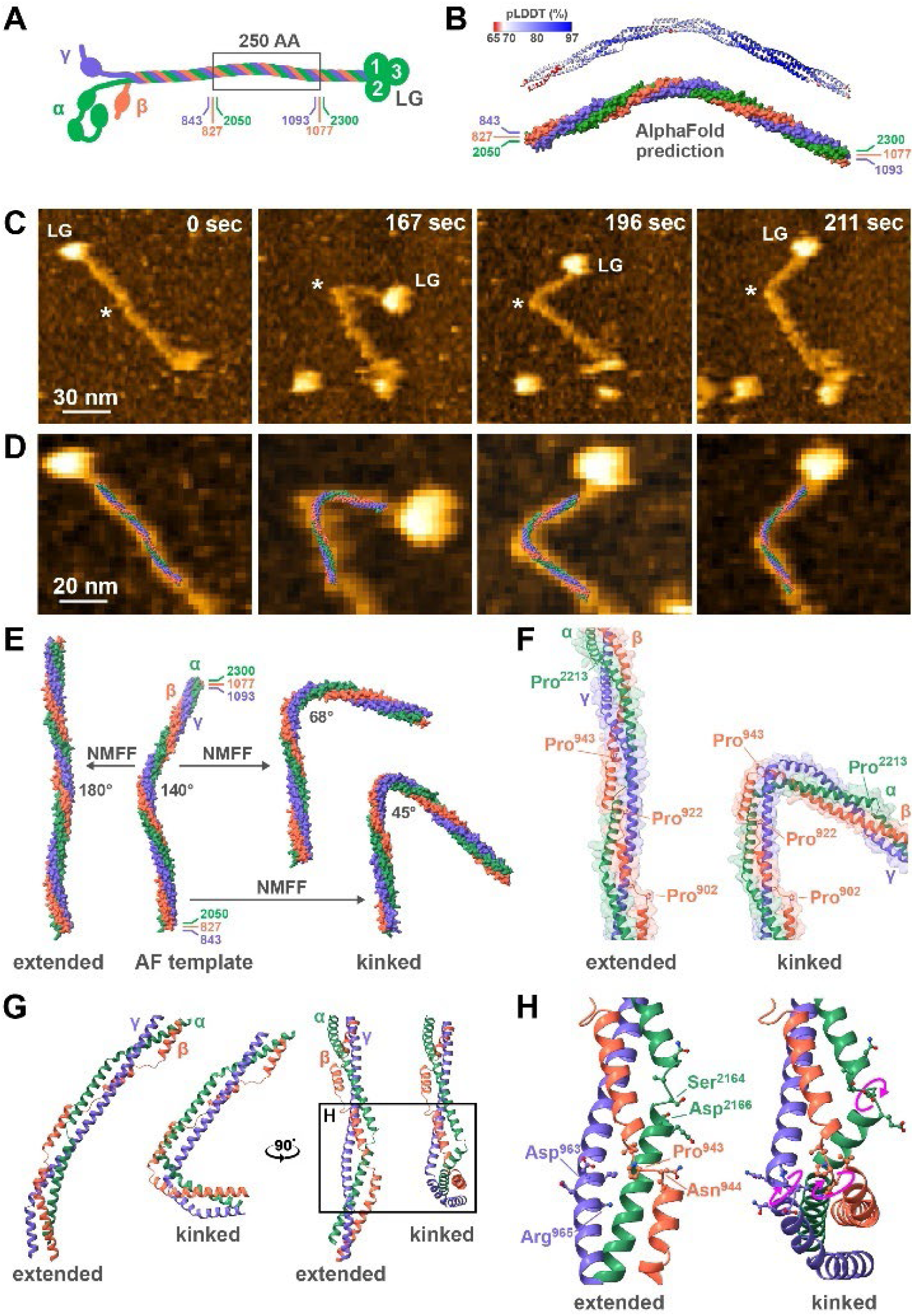
Elucidating dynamic coiled-coil hinge architecture. **(A)** Schematic depiction of the laminin-332 molecule indicating a central region (black box) of the coiled-coil domain spanning 250 amino acids (α3-chain 2050-2300, β3-chain 827-1077, γ2-chain 843-1093) used for AlphaFold structure prediction. **(B)** AlphaFold prediction of the hinge region of laminin-332 corresponding to the boxed region in **(A)** as a ribbon plot color-coded with the predicted local distance difference test (pLDDT, upper plot) and as a molecular surface plot (lower plot). **(C)** Stills from an HS-AFM time-lapse series depicting a single laminin-332 molecule in different coiled-coil conformations. The full range of the height scale is 7.5 nm. **(D)** Magnified views of the coiled-coil region displaying the superimposed AlphaFold coiled-coil segment from **(B)** after normal mode flexible fitting (NMFF) into the AFM topographies. **(E)** Representative coiled-coil structures in extended (left) or kinked (right) conformations obtained through NMFF of the AlphaFold template (middle) into AFM topographies. **(F)** Ribbon representation of the laminin-332 coiled-coil hinge region, superimposed onto a transparent surface plot, in the extended (left panel) and kinked (right panel) conformation. Proline residues associated with regions of local α-helix unwinding in the laminin α3- and β3-chain are indicated. **(G)** Side (left panel) and front (right panel) view of the laminin-332 hinge region in extended and kinked conformation, visualizing local α-, β-, and γ-chain unwinding during hinge kinking. **(H)** Magnified view of the hinge region. Several amino acid residues undergoing side chain rearrangement during kinking are highlighted, and pink curved arrows indicate the direction of side chain rotations.

AlphaFold structure template into any of the coiled-coil conformations present in the HS-AFM timelapse series (Fig. 3E and Movie 3), ranging from fully extended (∼180°) to acutely bent (∼35°) configurations. However, the fitted structures also revealed considerable conformational plasticity in the coiled-coil hinge region: while the hinge position remained static, the bending radius varied between frames, along with the bend angle (Movie 3). This indicated that sequences flanking the hinge (point of most acute coiled-coil bending) further modulate coiled-coil kinking. Analyzing the amino acid composition in this area revealed an unusual cluster of proline residues in the laminin β3-chain (Pro^902^, Pro^922^, Pro^943^) and α3-chain (Pro^2213^), coinciding with local α-helix unwinding (Fig. 3F). As α-helix breakers proline residues are sterically challenging for regular coiled-coil folding and therefore typically rare or absent in trimeric coiled-coil domains^53^. Comparing the extended and kinked coiled-coil conformations revealed that the short unwound stretch adjacent to Pro943 undergoes large conformational changes during kinking, suggesting that proline-induced α-helix unwinding may locally destabilize the coiled-coil domain and introduce flexible elements permitting coiled-coil bending. The β3-chain localizes to the outside of the hinge and may therefore need to undergo the most extensive conformational changes, including helix stretching, to facilitate coiled-coil bending. The array of regularly spaced proline residues separated by short α-helical segments in the β3-chain could provide the required molecular flexibility during coiled-coil bending, for instance through a “piston movement”-like translocation of the interjacent α-helical segments, as has been previously suggested as a mechanism for coiled-coil bending in other proteins^54, 55^. The molecular hinge displayed additional non-proline associated helix interruptions undergoing large conformation changes during kinking in the α-chain (Ser^2164^ - Asp^2166^) and γ-chain (Val^962^ - Arg^965^). Within these non-helical stretches, some residues displayed large side change rotations (α-chain: Asp^2166^, β-chain: Asn^944^, γ-chain: Arg^965^) during kinking (Fig. 3H). Thus, the staggered short stretches of unwound helix in the α-, β-, and γ-chains may define a precisely localized flexure point, while additional flexible elements placed farther away on either side of the hinge could provide additional flexibility, further modulating coiled-coil flexing and bending radius.

### Coiled-coil kinking directs elastase digestion towards producing the LM332E8 fragment

After characterizing hinge architecture and dynamics, we explored potential physiological roles of coiled-coil kinking and considered several possibilities, including regulation of laminin head/tail interactions, binding to other proteins, or susceptibility of laminin to proteolytic digestion. Interactions between the C-terminal LG domains and the laminin short arms have been proposed in the past, and severe coiled-coil bending could facilitate such intramolecular interactions. However, although coiled-coil bending typically reduced the distance between the C-terminal LG domains and the short arm cluster, we rarely observed direct contact between the terminal domains even in the most strongly bent molecules (kinking <30°) and could not identify the formation of stable intramolecular interactions in kinked molecules. In contrast to the laminin short arms and the LG domain cluster, which contain various protein interaction domains, the only known binding partner for the laminin long arm in or near the hinge region is the proteoglycan agrin^56^. Agrin binds to a central region in the laminin-111 coiled-coil domain, but critical residues in the γ1-chain mediating this interaction are absent in the γ2-chain, and agrin therefore does not bind laminin-332^57^. We instead investigated whether conformational changes could affect protease-mediated digestion of the laminin-332 coiled-coil domain, focusing on the serine endopeptidase elastase. Treatment with porcine pancreatic elastase (elastase-1, ELA1) cleaves laminin-111 into several well-characterized fragments, including the E8 fragment^52, 58, 59^. The E8 fragment consists of the C-terminal half of the long arm coiled-coil domain (∼35nm) and the subsequent LG domains and contains the integrin binding site at the coiled-coil domain/LG cluster interface^60^. Previous electron microscopy^59^ and biochemistry studies^52^ located the elastase cutting site in laminin-111 to a position roughly in the middle of the long arm, equivalent to the hinge region in laminin-332, raising the possibility that laminin-332 coiled-coil flexing could regulate elastase cutting in or near the molecular hinge. Visualizing conformation-dependent effects of protease digestion on individual laminin molecules requires fast imaging methods able to visualize the sample under physiological conditions, with single-molecule resolution and in real-time. Its high spatial and temporal resolution makes fast AFM scanning a unique tool to visualize such enzymatic activity on the single-molecule level^61–63^. To test if coiled-coil kinking affects coiled-coil cutting in laminin-332, we continuously imaged laminin-332 molecules by HS-AFM and then added a small volume (10 μl) of a porcine pancreatic elastase solution (1U/ml) directly into the fluid chamber (total volume 100 μl) of the HS-AFM. After several minutes the added enzyme had diffused into the imaging area and we started to observe elastase-mediated digestion of laminin-332, characterized by the cleavage of full-length laminin-332 into smaller fragments. This effect was elastase-dependent, since laminin-332 molecules imaged without elastase never displayed fragmentation even when imaged for extended periods (not shown). In these experiments we detected marked differences in the cleavage pattern depending on the laminin coiled-coil conformation. Laminin-332 molecules in the kinked conformation were cut first at or near the kink position (Fig. 4A, Movie 4), releasing a C-terminal coiled-coil segment attached to the LG1-3 domains (Fig. 4B). This fragment was morphologically indistinguishable from recombinant laminin-332 E8 fragment (Fig. 4C), in agreement with the previously reported site-specific cleavage of laminin by porcine pancreatic elastase into the E8 fragment. The N-terminal part of the coiled-coil was then often further digested, until only the short arm star remained (Movie 4), while the E8 fragment remained stable and was not further digested. The positions of the initial elastase cutting site (Fig. 4F) displayed excellent overlap with the range of hinge positions determined earlier (Fig. 2D), demonstrating that elastase cutting at or near the hinge in kinked molecules.

**Figure 4.**
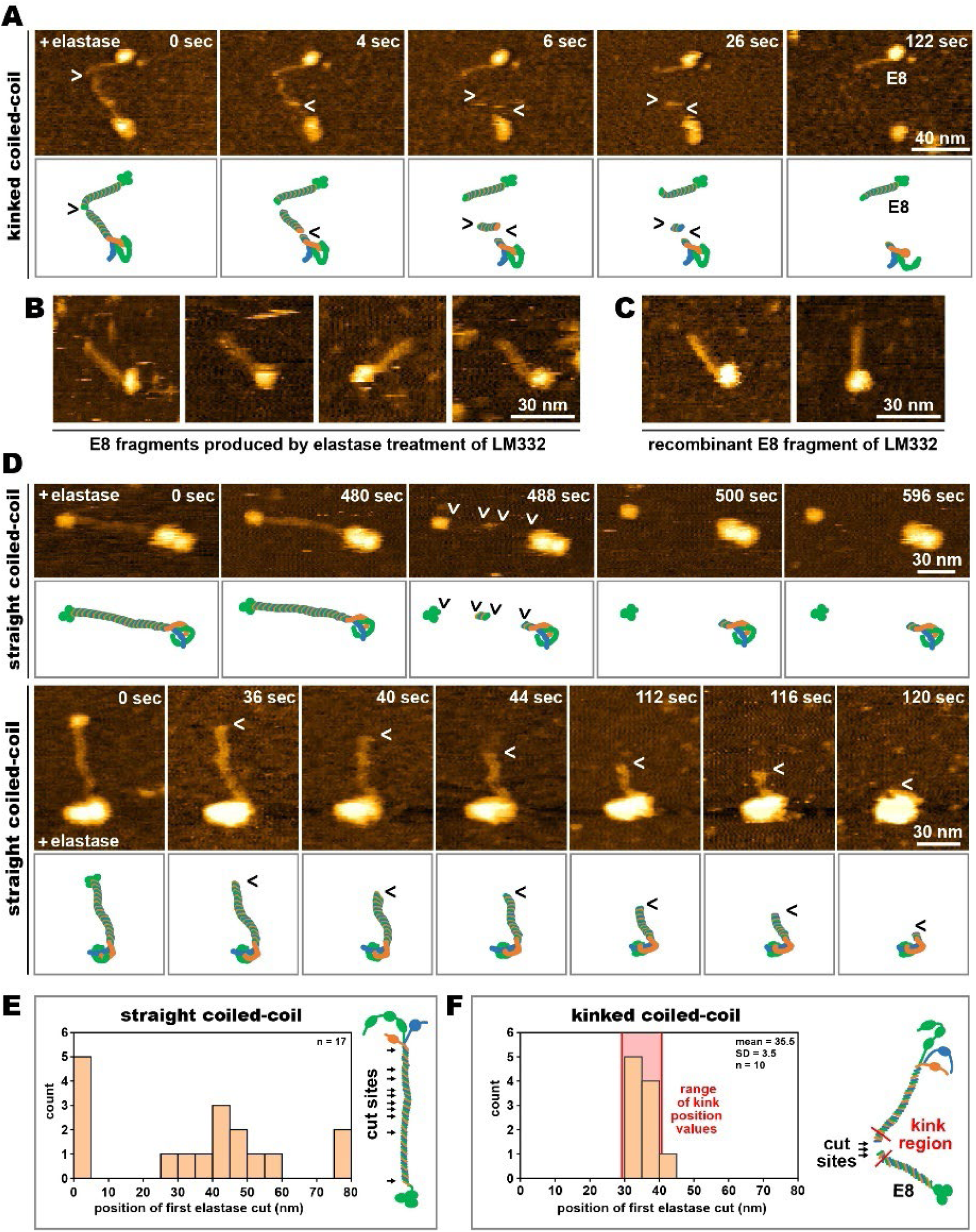
Visualizing coiled-coil conformation-dependent cleavage of laminin-332 by elastase. **(A)** HS-AFM time-lapse series visualizing elastase-mediated cleavage of a laminin-332 molecule in the kinked coiled-coil conformation (upper row) and corresponding schematic depictions of the digestion sequence (lower row). Arrowheads denote cleavage sites. The full range of the AFM height scale is 10 nm. **(B)** Representative images of E8 fragments produced by elastase digestion of laminin-332 molecules in the kinked coiled-coil conformation. The full range of the AFM height scale is 7 nm. **(C)** Representative AFM images of recombinant laminin-332 E8 fragments. The full range of the AFM height scale is 7 nm. **(D)** Two HS-AFM time-lapse series visualizing elastase-mediated cleavage of a laminin-332 molecule in the straight coiled-coil conformation (upper rows) and corresponding schematic depictions of the digestion sequence (lower row). Arrowheads denote cleavage sites. The full range of the AFM height scale is 10 nm. **(E)** Distribution of initial cleave location on molecules with straight (kink angle >126°) coiled-coil conformation (left panel) and schematic indication of cut locations within the laminin-332 coiled-coil (right panel). **(F)** Distribution of initial cleave location on molecules with bent (kink angle <126°) coiled-coil conformation (left panel) and schematic indication of cut locations within the laminin-332 coiled-coil (right panel).

In contrast to kinked molecules, laminin-332 molecules in extended conformations (bending angle >126°) were first cut in various positions along the coiled-coil domain, with enhanced frequencies near the beginning and end of the coiled-coil domain (Fig. 4D and 4E, Movies 4 and 5). In some cases, we observed spontaneous excision of the entire coiled-coil due to simultaneous cuts, or cuts happening in rapid succession, at the N- and C-termini of the coiled-coil (Fig. 4D). In other cases, there was successive shortening of the coiled-coil domain starting from the C-terminal end and progressing all the way to the point of short arm branching, consistent with a step-like pattern of elastase digestion (Fig. 4D and Movie 5). Regardless of the cleavage pattern and sequence (spontaneous excision or stepwise shortening), extended coiled-coil domains were ultimately fully removed from the laminin molecule, and E8 fragments were never generated. Together, these results indicate a conformation-dependent mechanism regulating laminin-332 digestion by elastase, with coiled-coil kinking guiding the initial elastase cut towards the hinge position and generation of the E8 fragment. The relation between coiled-coil kinking and elastase cutting is further supported by HS-AFM movies demonstrating extreme coiled-coil kinking (bending angle <30°) immediately prior to elastase cutting in the hinge region (Fig. 5A and B, Movie 6). Interestingly, such extreme kink angles <30° were never observed in absence of elastase (see Fig. 2C), suggesting that elastase binding promoted coiled-coil bending beyond the natural range driven by thermodynamic fluctuations in the elastase-free system. In some cases, apparent defects in the hinge region became visible just before and during extreme hinge kinking and prior to E8 fragment release (Fig. 5B and Movie 6). Possibly, one or two of the laminin chains are cut first, thereby decreasing bending strain and increasing hinge flexibility, until all three laminin chains are severed and the E8 fragment is released.

**Figure 5.**
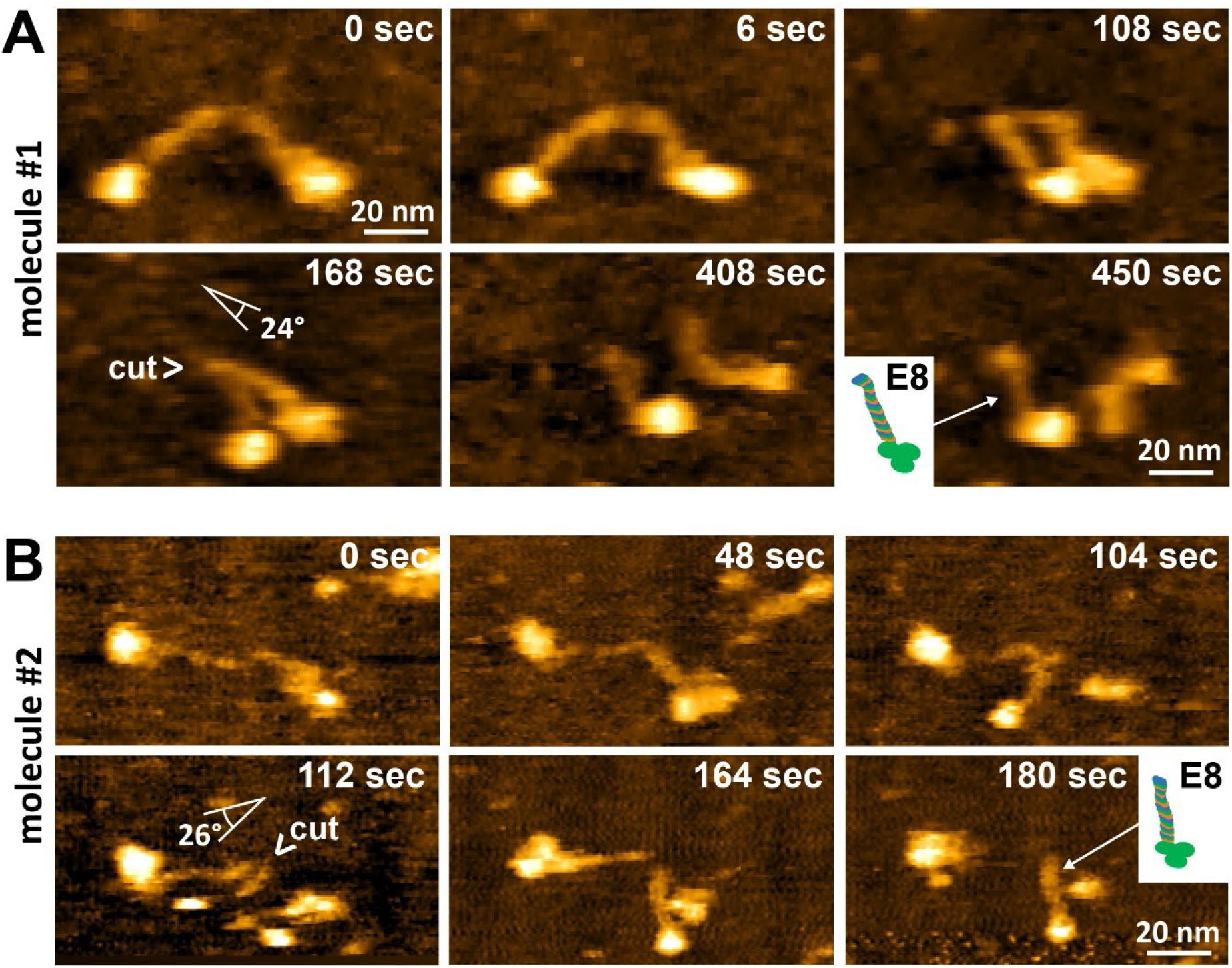
Acute coiled-coil kinking precedes laminin-332 hinge cutting by elastase. **(A)** and **(B)** stills from HS-AFM timelapse series of two laminin-332 molecules displaying acute coiled-coil kinking (24° and 26°) just prior to elastase mediated-hinge cutting and E8 fragment release. The full range of the AFM height scale is 10 nm.

**Figure 6.**
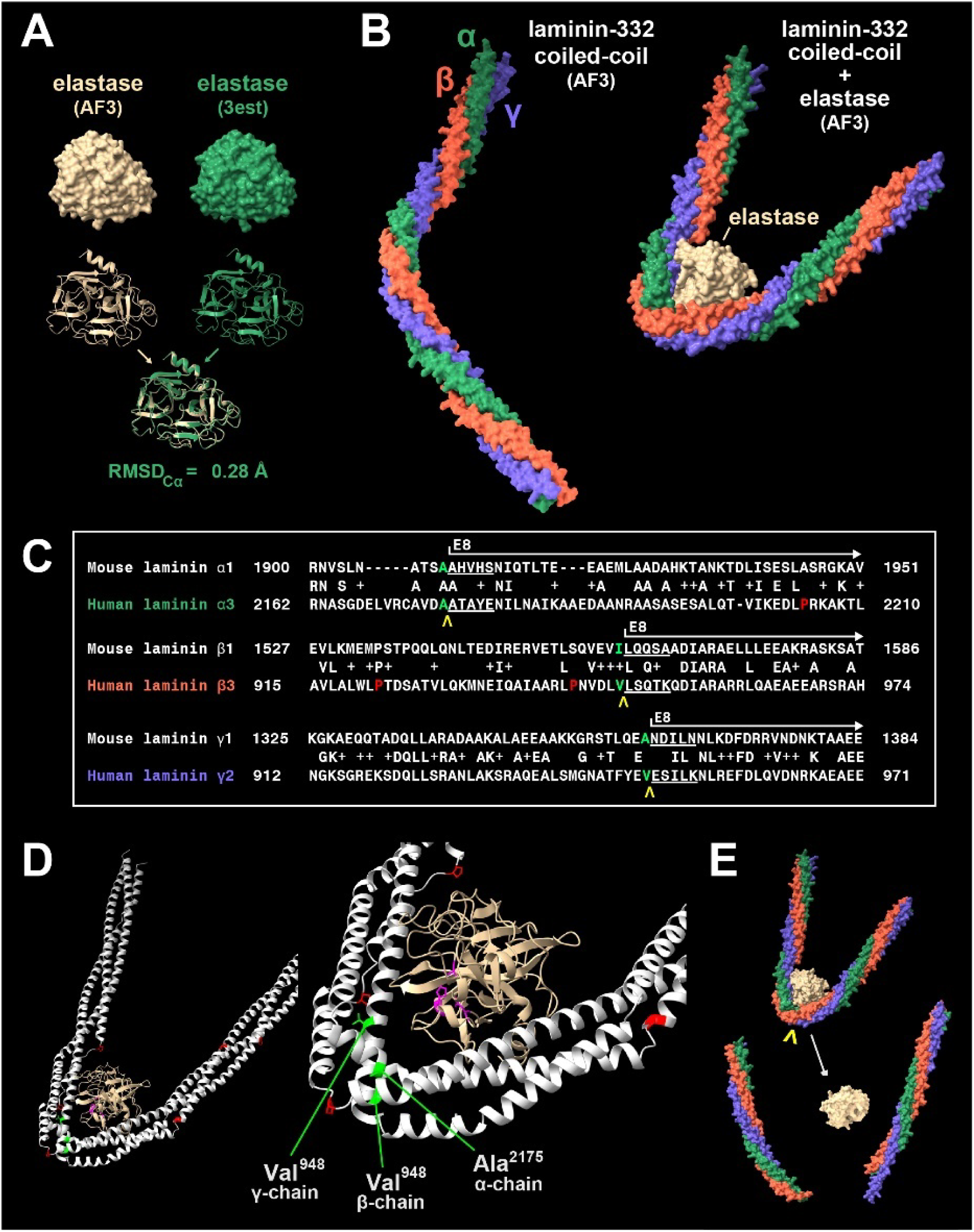
Elastase binding correlates with laminin-332 coiled-coil domain kinking. **(A)** Alphafold3 prediction (tan color) and crystal structure (3est, green) of porcine pancreatic elastase in space-fill and ribbon plots and overlay of both structures below. **(B)** Alphafold3 prediction of the central laminin-332 coiled-coil domain containing the molecular hinge alone (left panel) and in combination with elastase. **(C)** Blast protein sequence alignment of mouse laminin-111 and human laminin-332 across the elastase cutting site. The start of the E8 fragment in laminin-111 is indicated according to Deutzmann et al.^52^. The known (laminin-111) and putative (laminin-332) elastase cutting sites are indicated by yellow arrowheads, the N-terminal amino acid residues (P1) are in green. **(D)** Overview (left) and zoom-in (right) views of the laminin-332/elastase complex after molecular docking refinement showing the active site of elastase (cyan) in juxtaposition with the putative cleavage site in laminin-332 (P1 residues in green). Proline residues at sites of local coiled-coil unwinding plotted in red. **(E)** Space fill representation of the laminin/elastase complex before and after cleavage.

### Elastase binds inside the laminin-332 hinge and promotes coiled-coil kinking

To further investigate the relationship between elastase binding and coiled-coil bending, we assembled virtual elastase/laminin complexes in AlphaFold3. For this, we first confirmed accurate prediction of porcine pancreatic elastase structure by AlphaFold (Fig. 4A). Indeed, the AlphaFold structure was largely identical (RMSD_Cα_ = 0.23Å) to a crystal structure of the same enzyme^64^. As shown earlier (Fig. 3B), modelling of the laminin hinge region alone in AlphaFold yields a segment with moderate bending angle of ∼140°. However, inclusion of the elastase in the simulation predicted strongly increased coiled-coil bending around the elastase molecule binding to the inside turn of the hinge (Fig. 6B). The sharply decreased bending angle (∼25°) closely mirrored the low bending angles (24-26°) immediately before elastase cutting measured in the HS-AFM experiments (Fig. 5A and B). The AlphaFold rank 1 to 5 models all showed similar coiled-coil bending around elastase bound to the inside of the hinge, but slightly different elastase orientations (Suppl Fig. S2), suggesting some plasticity of the laminin/elastase interaction. These findings are consistent with a model in which elastase binding to the laminin hinge region stabilizes and augments the kinked conformation of the coiled-coil domain. In turn, the additional binding interfaces provided by the coiled-coil regions flanking the hinge may then trap the elastase molecule inside the laminin hinge, allowing time for optimization of the elastase orientation until its active site is directed towards targets sites in the hinge. While E8 digestion sites of the three laminin chains have not been determined in human laminin-332, they have been identified in mouse laminin-111 by peptide sequencing^52^. Aligning the laminin-332 hinge region with the corresponding sequences in laminin-111 using Protein BLAST revealed considerable sequence conservation of all three laminin chains in this region (Fig. 6C). Determining the laminin-332 residues corresponding to the known elastase cutting sites (P1 positions) in laminin-111 identified putative elastase cutting sites in the three laminin-322 chains (α3-chain: Ala^2175^, β3-chain: Val^948^, γ2-chain: Val^948^), which all localized directly within (α- and β-chain) or close to (γ-chain) the laminin-332 hinge (Fig. 6D). The localization of the putative α- and β-chain P1 residues to the sites of most acute kinking further supported a link between coiled-coil kinking and site-specific hinge cleavage by elastase. Moreover, all cleavage sites feature alanine (α3-chain) or valine (β3- and γ2-chain) residues in the P1 position, which are among the preferred residues for pancreatic elastase^65^. Due to a slight N-terminal offset of the γ2-chain site from the α3- and β3-chain sites inside the hinge, cutting at these putative sites would produce fragments with “sticky ends” with a ∼15 amino acid overhang of the γ2-chain in the E8 fragment (Fig. 6E). Indeed, some of the released E8 fragments displayed dynamic and frayed N-termini (Movies 4 and 6), although the expected overhang of ∼2.25 nm (15 x 0.15 nm, overhang retains α-helical fold) to ∼6 nm (15 x 0.4 nm, overhang fully extended) is close to the resolution limit of our AFM images and could not be resolved well. Together, these results suggest that elastase binding to the laminin-332 coiled-coil requires or induces extreme coiled-coil bending, stabilizing the elastase/laminin complex within the hinge, and directing elastase activity towards specific target sites in the hinge to produce the E8 fragment (Fig. 6E).

### Visualizing transient elastase/laminin complexes

To further clarify whether extreme coiled-coil kinking precedes and promotes elastase binding, or whether it is induced by elastase binding as suggested by the AlphaFold complex model, we attempted to visualize laminin/elastase complex formation and laminin digestion in real-time by HS-AFM. Typically, elastase-substrate complexes in solution are short-lived with durations of the catalytic event on the millisecond scale, but their lifetime may extend significantly on insoluble substrates, such as elastase fibrils^66^. Immobilization of the laminin-332 molecules on APTES-modified mica likely also lengthened the enzyme/substrate lifetime. Nevertheless, given the limited temporal resolution of HS-AFM (in our case 1 to 4 sec/frame), we did not usually observe the formation of the enzyme/substrate complex formation prior or during laminin cutting. However, in rare cases (3 out of 34 HS-AFM movies depicting laminin digestion), we detected potential elastase binding to the inside of the coiled-coil hinge in moderate bending states (70-90°, Fig. 7C and D and Movie 7). The experimental AFM image of the elastase/laminin complex closely resembled a simulated AFM image constructed from the Alphafold model of the complex (Fig. 7A and B). Within seconds of elastase binding, coiled-coil bending increased sharply down to angles of 20-30° (Fig. 7C). In this state the elastase molecule could not be discerned anymore, possibly because of burial within the acutely bent hinge. As observed before (Fig. 5A and B), shortly after transition into the strongly bent conformation, the laminin molecule was cut in the hinge positions and the E8 fragment was released. The sequence of events (elastase binding preceding sharp kinking) further suggests that elastase binding at or near the hinge region of laminin-332 stabilizes an acutely bent laminin conformation. In this way, the proteolytic activity of elastase could be directed towards the conserved cutting sites within the hinge. Ultimately, the kinking mechanism following elastase binding may ensure site-directed cutting of laminin to produce the biologically active E8 fragment, while exposure of laminin the extended coiled-coil conformation to elastase leads to spatially indiscriminate initial cutting and ultimately complete coiled-coil digestion (Fig. 8).

**Figure 7.**
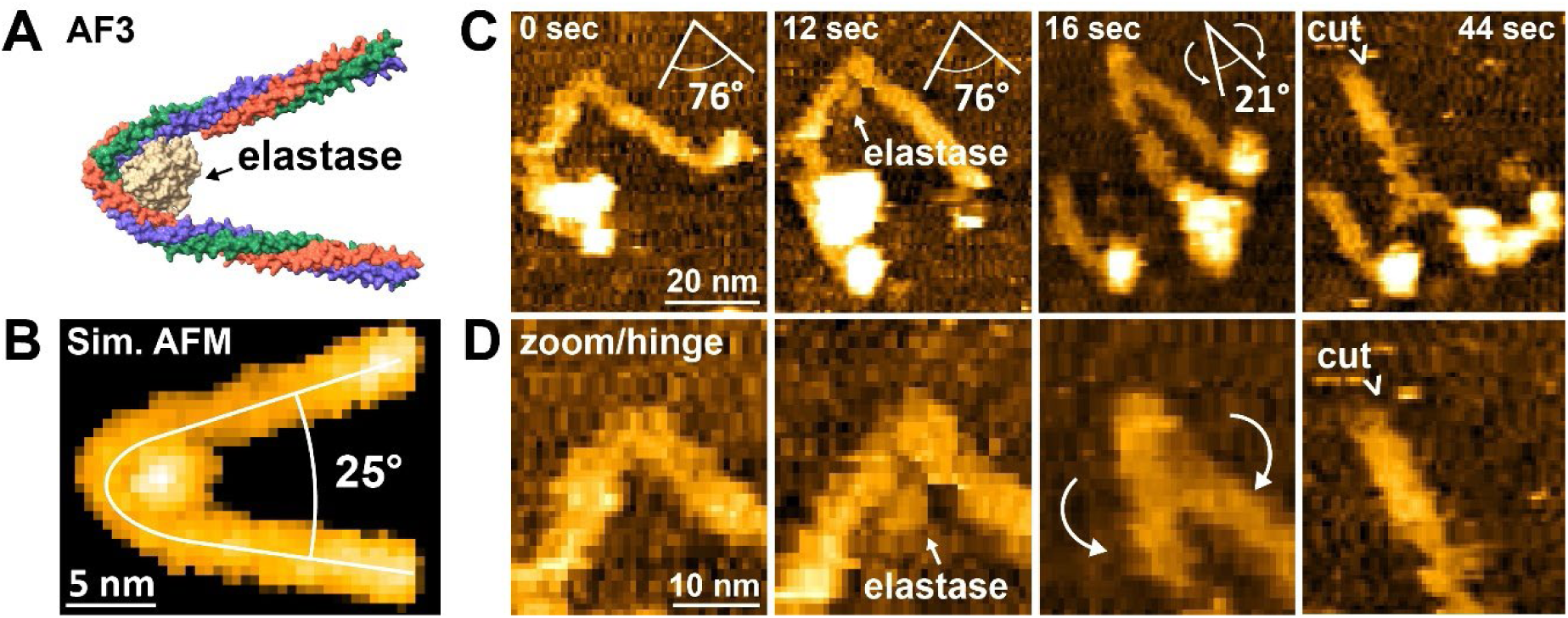
Visualizing transient elastase/laminin complexes during laminin cleavage. **(A)** AlphaFold3 model of the laminin-332/elastase complex and **(B)** the corresponding simulated AFM image. **(C)** Stills from a HS-AFM image series depicting putative elastase binding to a kinked (76°) coiled-coil domain, subsequent induction of acute kinking (21°) and hinge cleavage. The images also show a previously produced E8 fragment (lower left corners). **(D)** Zoomed-in views of the HS-AFM series shown in **(C)**.

**Figure 8.**
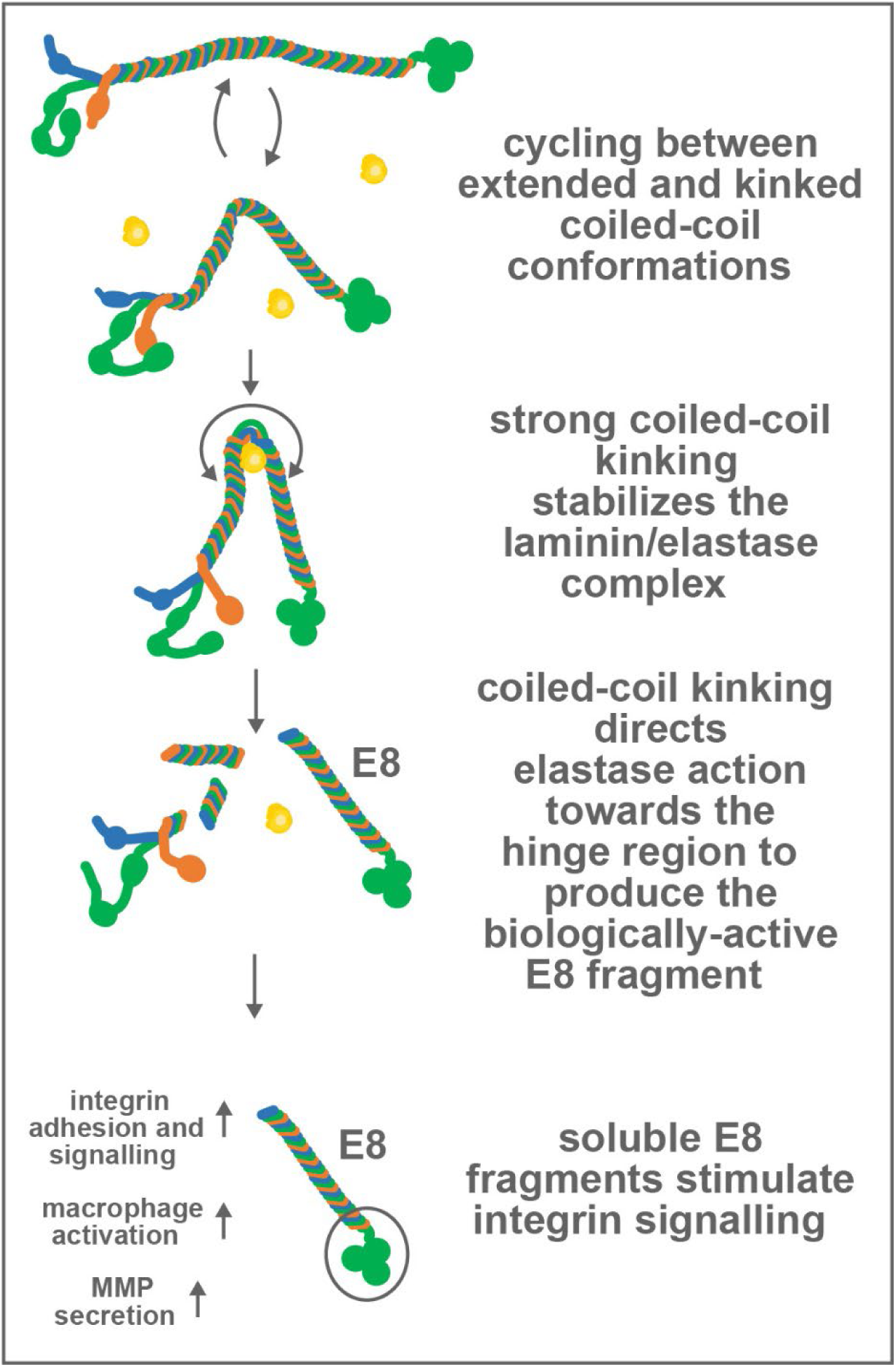
Model of conformation-dependent laminin-332 cleavage by elastase. Laminin-332 molecules constantly cycle between extended and kinked conformations. Kinking may stabilize transient elastase/laminin complexes by providing additional binding interfaces between elastase and the coiled-coil regions flanking the hinge. Severe kinking may then direct elastase activity towards a cutting site in the molecular hinge, while local coiled-coil unwinding may make this site more accessible to the protease. Reliable cutting in the hinge region ensures generation of the biologically active E8 fragment, which can then diffuse away from the original location of the laminin molecule in the basal membrane to stimulate integrins and other receptors for activating macrophages or other tissue-embedded cells.

## Discussion

### A new function for the laminin coiled-coil domain

In this study we identified a novel conformational control mechanism directing elastase cutting to a flexible hinge region in the laminin coiled-coil for producing the biologically active E8 fragment. These findings expand our understanding of the structure/function relationship of the laminin long arm, which has so far been best understood as a flexible molecular spacer separating the various protein interaction domains within the short arms from the integrin binding site at the C-terminal coiled-coil interface with the LG domain cluster. Elastase-mediated cleavage of the laminin-332 coiled-coil domain differed fundamentally depending on the coiled-coil conformation: while molecules in the extended conformation were cut in various positions along the coiled-coil, kinked molecules were exclusively cut first within the molecular hinge, producing the E8 fragment. We did not observe further cleavage of the E8 fragment once it was released from the kinked laminin-332 molecules, suggesting that the C-terminal coiled-coil region contained in this fragment is protected from further elastase digestion by an unknown mechanism. In contrast, extended molecules were frequently cut within the C-terminal half of the coiled-coil and sometimes even displayed successive trimming starting from the C-terminal end of the coiled-coil just adjacent to the LG domain cluster and progressing towards the N-terminal coiled-coil half, ultimately leading to complete coiled-coil degradation. *In vitro*, the processing of laminin by elastase can be guided into specific fragments (including E8) through limited proteolysis and careful choice of mild reaction conditions, such as long reaction times at low temperature (4° C) and low enzyme to substrate ratio^38, 59, 67^. In native tissues, other regulatory mechanisms are likely required to control laminin fragmentation. Guiding elastase cutting to the hinge region through coiled-coil kinking may be such a mechanism to ensure reliable generation of the biologically-active soluble E8 fragment, instead of full laminin degradation into non-functional small fragments.

Regardless of coiled-coil conformation, laminin molecules were frequently also cut at the N-terminal coiled-coil end, where a well-characterized neutrophil elastase target site exists in the γ2-chain of laminin-332 close to the base of the short arm aster^27^. In laminin-111 cutting by pancreatic elastase in this region also releases an E1-4 fragment containing the short arms^59^. However, extended molecules were often cut at this site first, while in kinked molecules such cutting occurred only after initial cleavage of the hinge. This suggests a mechanism by which coiled-coil kinking either delays cutting at the terminal sites, or more likely, accelerates cutting at the hinge. In kinked molecules the coiled-coil regions flanking the hinge may provide additional stabilizing binding interfaces for the elastase molecule, promoting successful enzyme/substrate complex formation and consequently accelerating cleavage at the hinge site. The twofold softening of the hinge after switching into the kinked conformation we determined may provide additional molecular flexibility for establishing a productive enzyme/substrate complex^68^, while partial coiled-coil unwinding in the hinge could ease access of elastase to the target cleavage sites. Preferential cutting in the hinge region may therefore be the result of different cleavage site kinetics, making it less likely to observe rarer or slower non-hinge cutting events before cutting at the favorable hinge site. In agreement with this interpretation, in HS-AFM movies showing extended and kinked molecule side-by-side, kinked molecules were typically cut first, while extended molecules displayed delayed cleavage (see for example Movie 4). However, HS-AFM-based single-molecule assay cannot assess enzymatic kinetics quantitatively, as the time between elastase addition into the AFM sample chamber, diffusion of the protease molecules into the AFM imaging area cannot be controlled precisely.

While our results demonstrate a clear relationship between coiled-coiled kinking and elastase cutting in *in vitro* experiments, it is unknown if laminin-332 exists in different coiled-coil conformations in native basement membranes, and if so, how coiled-coil kinking could be regulated. Laminin-332 is the central linker protein in the dermal-epidermal junction, which constitutes the critical mechanical interface between the dermal and epidermal skin layers and experiences large tensile and shear forces^69^. At the dermal-epidermal junction, the laminin-332 short arms interacts with collagen VII and other matrix components, while C-terminal binding sites mediate binding to cell surface receptors, including α_4_β_6_ integrin and syndecan-1 at hemidesmosomes^70^. The interjacent coiled-coil domain may therefore be subject to tensile force acting across the laminin-332 molecule. Considering the spring-like behavior of the coiled-coil domain^71^, variations in tensile forces across the dermal-epidermal junction could modulate laminin coiled-coil conformation and in this way result in biomechanical regulation of laminin cleavage and E8 fragment release. Interestingly, mechanical force also increases the availability of binding sites for pancreatic elastase in lung elastin and promotes elastase-mediated digestion^72^, suggesting a common mechanosensitive digestion mechanism of different matrix components by elastase.

### Directing elastase activity towards producing the biologically active E8 fragment

In this study we used porcine pancreatic elastase based on its well-characterized ability to cleave different laminin isoforms *in vitro*^59, 73^. Pancreatic elastase is a serine protease involved in digestive proteolysis of elastin and other dietary proteins in many animals^74^. However, despite its name the orthologous human pancreatic elastase (ELA-1) is transcriptionally silenced in human pancreas and plays no role in food digestion^75, 76^, which is instead carried out by chymotrypsin-like elastase family members 3 A and 3B (CELA3A, CELA3B)^77^. Instead, ELA-1 is expressed in the basal cell layer of the human epidermis^78^. The physiological relevance of the identified conformational control of laminin-332 digestion by elastase could therefore relate to tissue functions other than the digestion of dietary proteins. Given the crucial importance of laminin-332 for maintaining firm contact between the dermal and epidermal layers^69^, ELA-1-mediated cleavage of laminin-332 could for instance contribute to keratinocyte detachment from the basement membrane and subsequent movement to upper layers, and laminin kinking may contribute to regulating this process. Moreover, laminin processing by neutrophil elastase (NE, ELA-2), which is structurally and functionally related to ELA-1^79^ and cleaves laminin into similar fragments albeit significantly more efficiently^38^, has been linked to different pathologies. For instance, inflammation-induced neutrophil activation in the lung can lead to aberrant NE release, laminin-111 cleavage and the awakening of dormant cancer cells via stimulation of α3β1 integrin signaling^80^. Crucial in these cases may be the generation of soluble laminin fragments containing the integrin binding site that can diffuse away from the insoluble laminin layer within the basement membrane to stimulate integrin signaling in a wider tissue environment. E8 fragments of different laminin isoforms, including laminin-332, retain full integrin binding and stimulation capacity and are therefore candidate cleavage products in these scenarios. Recombinant E8 fragments are now widely used to promote cell adhesion, differentiation, and maintenance and expansion of pluripotent stem cells^81, 82^. Despite the proven effectiveness of recombinant E8 fragments in numerous *in vitro* applications, there are only few reports showing that E8 fragments are generated in native tissues^83^. The intrinsic conformational control mechanism in laminin-332 ensuring reliable release of the C-terminal E8 fragment identified in our study suggests that elastase cleavage is a specific and regulated process that could occur in physiological tissue environments.

### Possible kinking in other laminin isoforms

In our experiments we immobilized laminin-332 on APTES-modified mica, which provides stable sample attachment for HS-AFM scanning while maintaining sufficient molecular flexibility to monitor laminin dynamics. Under these immobilization conditions, only laminin-332 displays dynamic flexing around a sharply-defined coiled-coil hinge, whereas laminin-111 assumes a stable, S-shaped coiled-coil conformation^40^. Similar S-shaped, immobile coiled-coil domains were also observed for recombinant laminin-211, laminin-421, laminin-511, and laminin-521 (our observations). However, previous rotary shadow electron microscopy images of laminin-111 also show a distinct kink in the center of the coiled-coil and a variety of kink angles^38^, consistent with dynamic flexing around a defined molecular hinge. In fact, based on the approximate overlap of the laminin-111 coiled-coil kink position in EM images and the known elastase cutting site in this isoform, a link between coiled-coil flexibility and enhanced protease susceptibility has been proposed repeatedly^34, 38, 84^. However, a direct link could not be established at the time in absence of imaging techniques able to visualize dynamic laminin coiled-coil structure and elastase digestion simultaneously in individual molecules. Moreover, based on protein sequence alignment, the laminin-332 residues corresponding to the elastase E8 cutting sites in laminin-111^52^ localize precisely in the laminin-332 hinge, which in turn suggests a similar coincidence of hinge position and elastase cutting site in laminin-111. It is therefore possible that in solution or under different immobilization protocols, laminin-111 digestion by elastase may be similarly controlled by coiled-coil kinking. NE-mediated laminin-111 digestion plays an important role during inflammation^80^, and kinking could ensure generation of integrin-activating E8 fragments. Nevertheless, the drastically different coiled-coil domains shapes and dynamics in surface-immobilized laminin-332 and laminin-111 suggest that elastase susceptibility of different laminin subtypes is fine-tuned by coiled-coil structure and mechanics. On the other hand, the central coiled-coil domain of heart laminin (laminin-221) is resistant to elastase digestion altogether^85^, indicating that cleavage at the E8 target site is controlled individually in different laminin isoforms. Lastly, laminins are heavily glycosylated, mainly by asparagine (N)-linked oligosaccharides^86, 87^. The majority of potential N-glycosylation sites reside in the coiled-coil domain^38^, and N-glycosylation has been proposed to protect laminin-332 from proteolysis^88^. Thus, susceptibility to elastase digestions could be further modified by different N-glycosylation patterns within the laminin family.

### Mechanisms controlling coiled-coil kinking

Coiled-coils are common structural features in many proteins and can sometime span up to hundreds of nanometers. Proteins with these extended coiled-coil stretches frequently contain highly flexible hinge regions, and bending around these flexure points can lead to drastic conformational changes affecting protein function. For instance, in its inactive form the dimeric coiled-coil domain of the motor protein myosin II is backfolded, but phosphorylation of the myosin regulatory light chain induces heavy chain coiled-coil straightening, enabling myosin II self-assembly into thick contractile filament bundles^89^. Similarly, kinesin-1 motors assume a compact, folded and autoinhibited conformation through an “elbow” region within its coiled-coil domain, whereas activation requires coiled-coil straightening^90^. As another example, binding of the GTPase Rab5 to the tethering protein EEA1 induces a long-range conformational change from a rigid, extended to a flexible, collapsed state^91^, but the kinking mechanisms and precise hinge locations in EEA1 are still unknown. Bending around defined hinge elements is common but not limited to dimeric coiled-coil domains. For instance, cellular entry of the SARS-CoV-2 virus requires dramatic unfolding of a trimeric coiled-coil domain in the S2 subunit of the spike protein^92^. While coiled-coil kinking around precisely defined molecular hinges is therefore common, currently there is no satisfactory understanding which precise modifications to the heptad repeat sequence confer such highly localized flexibility. Deviations from the strict heptad repeat patters, such as “skips” (insertion of an extra residue) and “stammers” (insertion of a three residue discontinuity) are known to introduce bends and turns into coiled-coil domains^93^. In addition, zinc finger motives^94^ can introduce stable kinks in coiled-coils, However, these structural motives may introduce permanent kinks and bends in the coiled-coil and are therefore less suitable for enabling reversible hinge flexing. Additional molecular mechanisms must therefore exist that weaken the coiled-coil helix in precisely defined locations to facilitate reproducible and reversible coiled-coil unfolding/refolding. At the same time, the structural demands on these short flexible elements are high, as they must be able to support both regular linear coiled-coil structure, as well as sharply bent conformations while limiting wide-spread coiled-coil unraveling. Additional mechanisms have been implicated in facilitating reversible coiled-coil kinking, including local coiled-coil unwinding^95^, “piston movements” of α-helix segments^55, 96^, presence of asparagine residues in the first or fourth position of the heptad^97, 98^, and prolyl cis/trans isomerization^99^. The unusual series of regularly spaced proline residues in the hinge region of the laminin β3-chain suggest a functional link to laminin-332 coiled-coil bending. Proline residues introduce pronounced kinks into α-helices and thereby function as helix breakers^100, 101^. Furthermore, prolines typically also force the N-terminal three amino acids into a loop structure^102^. Such α-helix disruptions put steric constraints on regular coiled-coil supercoiling and consequently prolines are rare in trimeric coiled-coils, with a prevalence of <0.4% of total residues, compared to 6.3% in the overall proteome^53, 103^. Given their extreme rarity in trimeric coiled-coils, little is known about the potential role of prolines in regulating trimeric coiled-coil bending. In dimeric coiled-coils however, the presence of prolines can coincide with coiled-coil bending or increased flexibility. For instance, the flexible “elbow” loop in kinesin family member 5A (KIF5A), required for the transition between a compact, inactive and extended, active kinesin conformation contains a highly conserved proline residue^104^. Of note, all proline residues within the laminin-332 hinge region (β3-chain: Pro^902^, Pro^922^, Pro^943^, α3-chain: Pro^2213^) induce short N-terminal α-helix unwinding as expected. The staggered array of short α-helical segments interspersed by short proline-dependent unwound stretches could provide the necessary flexibility and chain extendibility for coiled-coil bending, akin to the corrugated hinge element of a flexible drinking straw. In future, detailed mutational studies will be necessary to elucidate the precise role of these proline residues for laminin-332 coiled-coil kinking and digestion.

Cryo-electron microscopy studies have generated atomistic structures of several dimeric coiled-coil hinges, typically in the bent conformation^105, 106^. This is possible if the bent conformation is stable, allowing for particle averaging, but generating structural models of dynamic hinges is still challenging. The introduction of advanced flexible fitting of molecular models into resolution limited AFM images has now greatly expanded the capability of HS-AFM for obtaining dynamic atomistic structures^42^. Besides deepening our understanding of the molecular mechanisms driving coiled-coil kinking, the obtained insight may also aid the development of artificial peptide-based nanostructures with defined and addressable mechanical properties for cell function regulation^107^. Coiled-coil domains are often incorporated into such designer nanostructures as interaction mediators or molecular spacers^108, 109^. Better understanding molecular mechanisms underlying coiled-coil flexibility will greatly benefit the rational design of macromolecular scaffolds with flexible linker elements permitting nanostructure flexing. Moreover, structural insight into coiled-coil hinge architecture is now aiding in the rational design of activating peptides able to stabilize extended coiled-coil conformations in native proteins^110^. By visualizing the full spectrum of intermediary conformations of flexing coil-coil hinges, HS-AFM in combination with molecular fitting algorithms can contribute important structural insight for the design of candidate peptide modulators and visualize their effect on coiled-coil bending in real time.

## Acknowledgments

This work was supported by the Ministry of Education, Culture, Sports, Science and Technology (MEXT, https://www.mext.go.jp), Japan, through the World Premier International Research Center (WPI) Initiative (HF, CB, RA and CMF), and by the Japan Science and Technology Agency (https://www.jst.go.jp) through CREST No. JPMJCR1762 (HF). This work was also supported by a WPI-NanoLSI Transdisciplinary Research Promotion Grant, Kanazawa University. LA was also supported by JST, the establishment of university fellowships towards the creation of science technology innovation, Grant number JPMJFS2116 and WPI NanoLSI, Kanazawa University.

## Author contributions

C.M.F. devised the study and wrote the manuscript. L.A. and C.M.F. performed HS-AFM experiments, processed AFM data, performed statistical analysis and prepared figures. H.F. and C.M.F. performed AlphaFold structure prediction experiments. H.F. and R.A. carried out flexible fitting and molecular docking experiments. C.B. and R.G. performed free energy calculations and lifetime estimations. All authors read, revised, and approved the manuscript before submission.

## Movie legends

**Movie 1. Cycling of laminin-332 between extended and kinked coiled-coil conformations.** HS-AFM movie recorded at 1 frame/per second showing dynamic cycling of laminin-332 between extended and kinked coiled-coil conformations. The full range of the height scale is 8 nm.

**Movie 2. Normal mode flexible fitting to reveal dynamic coiled-coil hinge architecture.** HS-AFM movie (left), superimposition of NMFF structures (middle), and NMFF structures alone (right). The full range of the height scale is 8 nm.

**Movie 3. Visualizing coiled-coil hinge dynamics with atomistic resolution.** Dynamic model obtained by NMFF of an AlphaFold3 prediction of the central coiled-coil domain area (250 AAs) containing the molecular hinge into resolution-limited AFM topographies. Laminin α-chain in green, β-chain in red, γ-chain in blue.

**Movie 4. Different elastase digestion patterns of extended and kinked laminin-332 molecules.** HS-AFM movie recorded at 0.5 frame/per second showing different elastase digestion patterns of laminin-332 molecules in extended or kinked conformations. Elastase had been added to the HS-AFM sample chamber 5 min before the start of recording. Cleavage of the kinked laminin molecule occurs in the hinge region and produces the E8 fragment. The full range of the height scale is 10 nm.

**Movie 5. Full coiled-coil domain degradation after elastase digestion of extended laminin-332 molecules.** Representative HS-AFM movies recorded at 0.25 frame/per second showing the effect of elastase on two representative laminin-332 molecules with extended coiled-coil conformation. Addition of elastase induces excision (left movie) or gradual (right movie) coiled-coil domain degradation. The full range of the height scale is 8 nm.

**Movie 6. Acute coiled-coil kinking precedes elastase mediate hinge cutting.** Two HS-AFM movies showing elastase-mediated cutting at the molecular hinge in the coiled-coil domain. Movies are paused to visualize increased kinking preceding the cleavage step in the hinge. The full range of the height scale is 9 nm.

**Movie 7. Visualizing a transient laminin-332/elastase complex before hinge cutting.** A HS-AFM movie (0.25 sec/frame) showing elastase binding to the laminin-332 coiled-coil hinge (movie briefly pauses). Afterwards, coiled-coil kinking increases and the coiled-coil domain is cleaved at the hinge location. The full range of the height scale is 8 nm.

