## Supplemental Information for "Coiled-coil domain kinking controls laminin-332 cleavage by elastase"

### *Coiled-coil kink lifetime estimation*

**Maximum-likelihood estimation of *D*.** The estimation of the transition rate between the two characteristic kink angles is based on the Langevin dynamics

$$\dot{\varphi}(t) = Dv(\varphi) + \sqrt{2D}\,\xi(t).$$

in which $v(\varphi) = d\log P_s(\varphi)/d\varphi$ is an abbreviation. The corresponding Fokker-Planck equation for the conditional probability density $P(\varphi, t|\varphi_0, t_0)$ to find a kink angle $\varphi$ at time $t$, given that the process started with an angle $\varphi_0$ at time $t_0$, is given by

$$\frac{\partial P(\varphi, t|\varphi_0, t_0)}{\partial t} = -D\frac{\partial}{\partial\varphi}\left[v(\varphi)P(\varphi, t|\varphi_0, t_0) - \frac{\partial P(\varphi, t|\varphi_0, t_0)}{\partial\varphi}\right].$$

Its short-time solution ($|t - t_0| \ll 1$) is a Gaussian (Risken, 1996) with mean $\mu = \varphi_0 + Dv(\varphi_0)(t - t_0)$ and variance $\sigma^2 = 2D(t - t_0)$:

$$P(\varphi, t|\varphi_0, t_0) \approx \frac{1}{\sqrt{4\pi D(t - t_0)}}\exp\left\{-\frac{[(\varphi - \varphi_0) - Dv(\varphi_0)(t - t_0)]^2}{4D(t - t_0)}\right\}.$$

Accordingly, the likelihood $\mathcal{L}_\phi(D)$ of the parameter $D$, given an observation of a time series of $N$ angle measurements $\phi = \{\varphi_1 = \varphi(t_1), \varphi_2 = \varphi(t_2), \dots, \varphi_N = \varphi(t_N)\}$, is determined by a product of Gaussians

$$\mathcal{L}_\phi(D) = P_s(\varphi_1)\prod_{n=2}^{N} P\left(\varphi_n, t_n|\varphi_{n-1}, t_{n-1}\right).$$

We estimate the parameter $D$ by maximizing the likelihood function $\mathcal{L}_\phi(\mathcal{D})$ with respect to this parameter: $\widehat{D}_{\mathrm{ML}} = \arg\max_D \mathcal{L}_\phi(D) \approx 0.12\,\mathrm{rad}^2/\mathrm{s}$. Note that the factor $P_s(\varphi_1)$, corresponding to the initial configuration, does not contribute to the likelihood as the stationary distribution is independent of $D$ which solely sets the timescale of the dynamics.

**Numerical solution of the Langevin dynamics.** We simulate the Langevin dynamics by discretizing time (numerical time step $\tau = 10^{-2}$ s), employing a Euler-Maruyama scheme:

$$\varphi(t + \tau) = \varphi(t) + \tau Dv(\varphi) + \sqrt{2D\tau}\,\sigma_t$$

with mutually independent random numbers $\sigma_t$ drawn from a standard normal distribution (Risken, 1996).

### *Reference*

Hannes Risken: The Fokker-Planck Equation. Methods of Solution and Applications (Springer, 1996).

***Supplemental Figure 1***

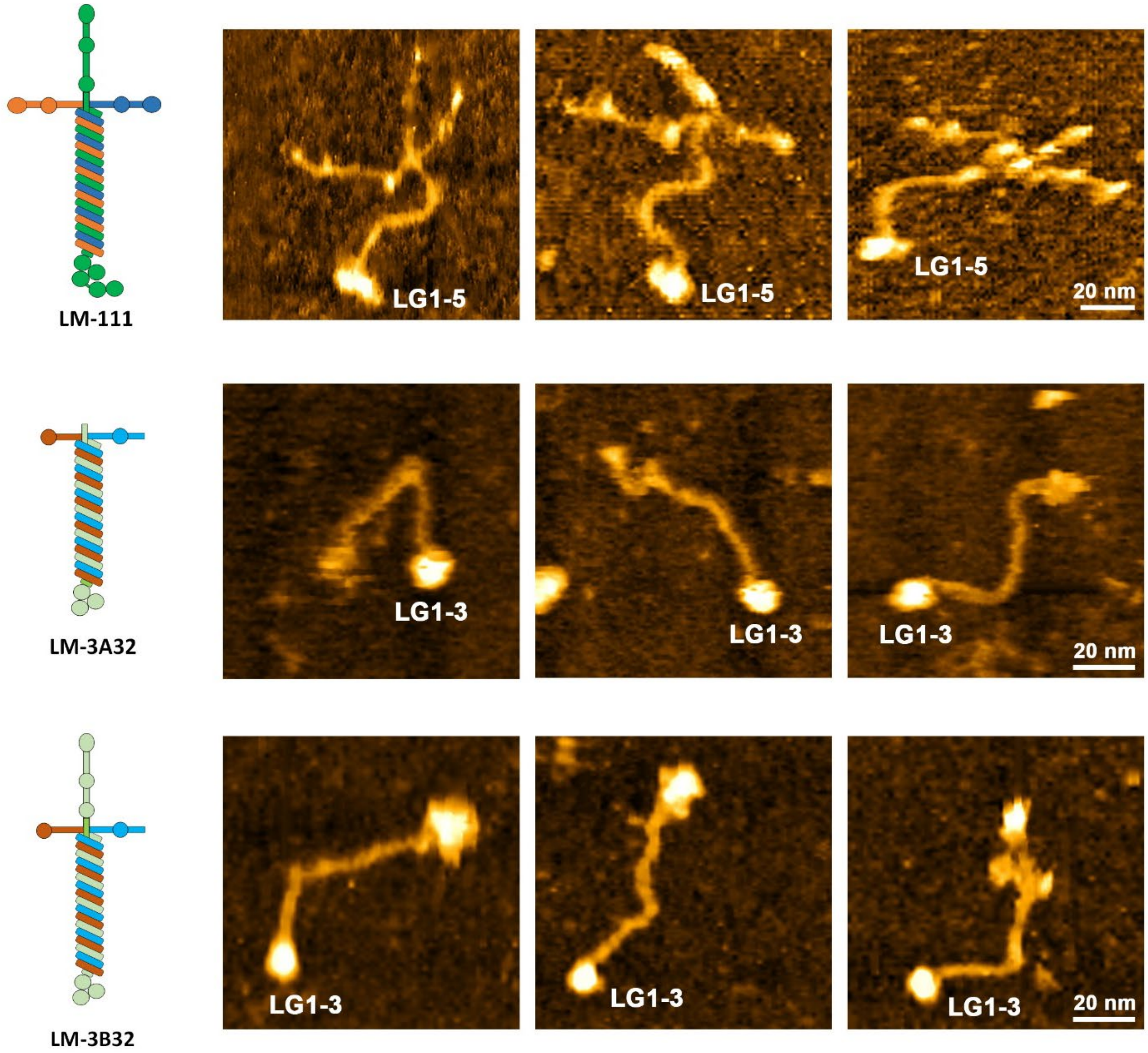


**Suppl. Fig. 1.** Representative HS-AFM image stills of recombinant Human laminin-111 (top), laminin-2A32 (middle), and laminin-3B32 visualizing different coiled-coil shapes. The full range of the height scale is 15 nm.

***Supplemental Figure 2***

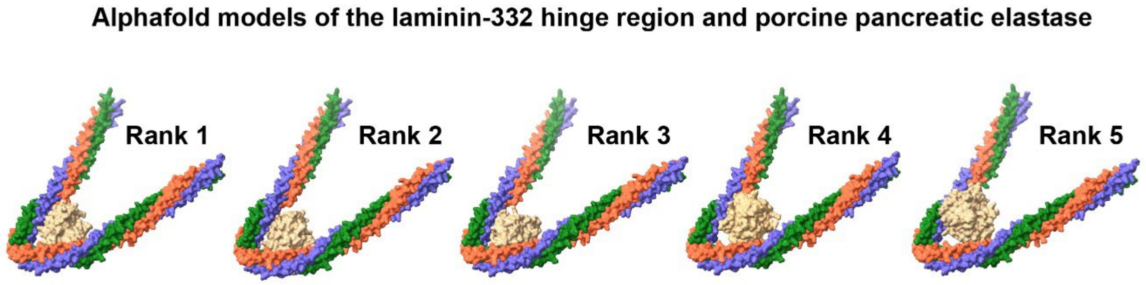


**Suppl. Fig. 2.** The rank 1-5 models generated by AlphaFold3 of the laminin/elastase complex. While laminin hinge kinking (hinge position, bending angle) is comparable in all models, elastase position and orientation varies slightly between the models.
